# Worldwide study of the taste of bitter medicines and their modifiers

**DOI:** 10.1101/2024.04.24.590957

**Authors:** Ha Nguyen, Cailu Lin, Katherine Bell, Amy Huang, Mackenzie Hannum, Vicente Ramirez, Carol Christensen, Nancy E. Rawson, Lauren Colquitt, Paul Domanico, Ivona Sasimovich, Riley Herriman, Paule Joseph, Oghogho Braimah, Danielle R. Reed

**Affiliations:** Monell Chemical Senses Center, Philadelphia PA, USA; Clinton Health Access Initiative, New York, NY, USA; National Institute of Alcohol Abuse and Alcoholism & National Institute of Nursing Research, Bethesda MD, USA; Countess of Chester Hospital, Chester, UK

**Keywords:** medicine, taste, bitterness, bitter reducer, genetic, sensory evaluation

## Abstract

The bitter taste of medicines hinders patient compliance, but not everyone experiences these difficulties because people worldwide differ in their bitterness perception. To better understand how people from diverse ancestries perceive medicines and taste modifiers, 338 adults, European and recent US and Canada immigrants from Asia, South Asia, and Africa, rated the bitterness intensity of taste solutions on a 100-point generalized visual analog scale and provided a saliva sample for genotyping. The taste solutions were five medicines, tenofovir alafenamide (TAF), moxifloxacin, praziquantel, amodiaquine, and propylthiouracil (PROP), and four other solutions, TAF mixed with sucralose (sweet, reduces bitterness) or 6-methylflavone (tasteless, reduces bitterness), sucralose alone, and sodium chloride alone. Bitterness ratings differed by ancestry for two of the five drugs (amodiaquine and PROP) and for TAF mixed with sucralose. Genetic analysis showed that people with variants in one bitter receptor variant gene (*TAS2R*38) reported PROP was more bitter than did those with a different variant (p= 7.6e-19) and that people with either an *RIMS2* or a *THSD4* genotype found sucralose more bitter than did others (p=2.6e-8, p=7.9e-11, resp.). Our findings may help guide the formulation of bad-tasting medicines to meet the needs of those most sensitive to them.

## 1. Introduction

Bitter taste likely evolved to deter poison ingestion [1], and bitter compounds are added to some products to prevent accidental poisonings [2]. However, this adaptive bitter avoidance is maladaptive when beneficial, even life-saving medicines are rejected by patients [3]. This rejection often creates a challenge of administering many types of medicine, especially liquid formulations, due to their unpleasant, often bitter taste [3, 4]. Furthermore, clinicians report that a drug’s poor palatability is a barrier to completing and adherence to treatment [5-7] and the non-adherence is high in low-resource settings such as African countries [8-11]. Therefore, those who formulate medicines use excipients (ingredients like sweeteners) that they can add to the active pharmaceutical ingredient (the drug) to reduce bitterness, but often these additions are only partly effective at improving taste. Thus, investigators search for new ingredients that might improve the taste of medicines, to optimize medicines, so that all who need them can easily take them.

The concept of optimizing implies there is one formulation that will work best for all people. But when food manufacturers optimize food products, they tailor their formulations to local tastes (e.g., for snacks like chips or sodas in a specific community), with more resources allocated to tailoring the product to geographical markets. For international markets, often there is no single ideal version of a food product. Medicines, unlike snack foods, are essential commodities, but it is less common to formulate for a specific market, especially because there is little scientific knowledge in the public domain to improve taste based on how people differ in their perceptions of medicines or taste modifiers. Such knowledge could be useful economically because, since a formulation that appeals to those with the most difficult palate (and thus most likely to fail to take their medicine because of its taste) would easily be accepted by those less sensitive, a universal formulation could be developed for medicines.

There are several examples of how the taste of medicines can differ markedly among people. For example, early human sensory studies of the bitter-tasting antimalarial drug quinine showed person-to-person differences [12], and like color-blindness, genetic differences at least partially account for some people tolerating its taste [13-15]. The most sensitive quinine genotype differs by ancestry, but ancestry is often overlooked in quinine sensory tests [16]. As another example, the thyroid medications methimazole [17] and propylthiouracil (PROP) [18] markedly differ in bitterness ratings owing to genetic variation. For methimazole, the insensitive variant is present in about half of humans worldwide [19], but it is concentrated in people in specific geographic regions [20] – a result foreshadowed by early anthropologists who studied the taste of similar chemicals like PROP and phenylthiocarbamide [21]. Experts in the area of drug formulation have considered the effects of age on drug palatability, but rarely ancestry [16, 22-24].

Therefore, we hypothesize not only that the study of bitter medicines will uncover large person-to-person differences but also that these differences could correlate with geographic ancestry and genotype.

To test this hypothesis, and to better understand the range of human taste experiences with medicines, we chose medicines that are important worldwide in the treatment of infectious diseases: tenofovir alafenamide (TAF), moxifloxacin HCl, praziquantel, and amodiaquine HCl. For comparison, we also included PROP, a medicine already well known to be experienced differently by people based on genotype and ancestry. We asked people to rate their bitterness and evaluated their ancestry. To capture the range of genotypes from people in Asia and Africa, we enlisted research participants who were recent US and Canada immigrants from these continents (who are underrepresented in human sensory studies), plus a comparison group of people of European ancestry. We collected saliva samples from each person to examine their genetics, to learn more about how and why people differ in taste responses to these key medicines.

Fundamental taste qualities other than bitterness also differ among people, although for sweetness, saltiness, and sourness the range is less extreme [25]. These differences are partially due to inherited genetic variation, so they too can also differ by ancestry, e.g., [26, 27]. Thus, although adding sweet, salty, or sour flavorings is a common strategy to make medicines more palatable, these too are unlikely to work well for all people. To find more effective taste modifiers for medicines, investigators have sought a new generation of tasteless bitter blockers that work at the taste receptor [28], but as with drugs and classic taste qualities, their effectiveness differs among people [29]. Therefore, we also studied two excipients that reduce bitterness, the artificial sweetener sucralose and the tasteless bitter blocker 6-methylflavone (6- MF), for their effect on responses to TAF, a drug previously tested with 6-MF for bitter blocker efficacy [30].

## 2. Materials and methods

### 2.1. Drug selection, sourcing, and preparation

We selected bitter drugs from among those commonly used to treat infectious diseases of public health importance common in resource-limited countries in populations where there is little data on bitterness perception and among whom compliance is critical to achieve current public health goals [11]. We filtered out drugs that were (a) likely to be discontinued in the near future or (b) structurally similar to others on the list. The final selection comprised tenofovir alafenamide (TAF), moxifloxacin HCl, praziquantel, and amodiaquine HCl. ***Table 1*** lists the diseases these drugs are indicated for and their solubility. For praziquantel, we chose the racemic form (see **Supplemental Table A**), which is less expensive and prescribed more often than the chiral S and less bitter R form [31, 32]. Propylthiouracil (PROP) served as a control drug, since genetic effects on its bitterness intensity ratings are well known [e.g., [18, 33]]. We also included two solutions of TAF mixed with an excipient: sucralose, an artificial sweetener known to suppress bitterness [34], or 6-methylflavone (6-MF), a bitter blocker, at least in some people, which is tasteless [30] (***Table 1***). Drugs were purchased from the vendors listed in **Supplemental Table B.** We dissolved each drug in deionized water, ethanol, or other diluent.

**Table 1.** Characteristics of taste stimuli used in this study.

| Compound | PubChem CID | Use | Solubility (mg/mL) | Diluent | Taste | Sample concentration (μM) |
| --- | --- | --- | --- | --- | --- | --- |
| Tenofovir alafenamide fumarate (TAF) | 9574768 | HIV medication | 4.7 [68] | Deionized water | Bitter [69, 70] | 200 μM |
| Moxifloxacin HCl | 152946 | Tuberculosis medication | 24 [71] | Deionized water | Bitter [72] | 300 μM |
| Praziquantel | 4891<sup>a</sup> | Schistosomiasis medication | 10 [32] | 3% ethanol | Bitter [31, 73] | 300 μM |
| Amodiaquine HCl | 2165 | Malaria medication | 2.0 [74] | Deionized water | Bitter [75] | 300 μM |
| Propylthiouracil (PROP) | 657298 | Hyperthyroidism medication | 1.2 [76] | Deionized water | Bitter [76] | 560 μM |
| Sucralose | 71485 | Food additive (sweetener) | 300 [77] | Deionized water | Sweet [77] | 2.5 mM |
| 6-Methylflavone (6-MF) | 689013 | Bitter receptor blocker<sup>b</sup> | # [78] | 2.5% ethanol + 0.5% Tween-80 | -<sup>c</sup> [78] | 500 μM |
| Sodium chloride | 5234 | Food ingredient | 36.0 [79] | Deionized water | Salty [36] | 256 mM |

<sup>a</sup>racemic; <sup>b</sup>highly soluble in DMSO, details unspecified; <sup>c</sup>rated as “barely detectable” or “weak”

### 2.2. Pilot taste testing

We conducted pilot tests to identify a concentration of each drug that gave mid-range intensity ratings of bitterness (**Supplemental Table C**). We avoided concentrations that were near the bottom or top of the intensity rating scale to avoid floor and ceiling effects, so we could better identify person-to-person differences. We prepared ascending concentrations of each stimulus, plus a fixed concentration of each excipient alone: sucralose (2.5 mM) and 6-MF (500 µM). We also tested a fixed concentration of sodium chloride (256 mM), because saltiness is familiar to most people and easy to detect at this concentration [35], and because sodium chloride is another excipient used to reduce bitterness [36].

We convened an experienced sensory panel of 11 participants (aged 24-72 years; 8 [73%] females; 9 [82%] Europeans, 1 [9%] Asian, and 1 [9%] Hispanic). They tasted and expectorated (spit out) ascending concentrations of each drug following our previously published testing procedures [33]. Between each sample, panelists rinsed their mouths with water twice.

Panelists also provided open-ended comments on any aspects of the sensory experience (e.g., burning or tingling).

### 2.3. Study participants

#### Participant eligibility and recruitment

English-speaking adults over 18 years old (no upper age limit), either biological sex, and who were first-, second-, or third-generation immigrants from Asia or Africa by self-identification, or who were of European ancestry were eligible. We used four methods to recruit participants: (1) posted paper flyers on the campus of the university and other bulletin boards; (2) advertised on social media, including Facebook, LinkedIn, Reddit, TikTok, and Instagram; (3) asked past participants to refer eligible people from their professional and personal networks; and (4) contacted professionals in Asia and Africa and asked them to recruit prospective participants who had recently moved from their home country to the US or Canada. Participants were screened for COVID-19 by a questionnaire and excluded for any signs of illness on the day of sensory testing.

#### Ethics approval

All procedures were approved by the Institutional Review Board at the University of Pennsylvania (protocol #701426, *Genetics of Taste and Oral Perception*). The sip-and-spit taste procedures used for active pharmaceutical ingredients of Federal Drug Administration (FDA)- and World Health Organization (WHO)-approved drugs and common sensory compounds confer no more risk than from a routine medical visit.

### 2.4. Taste testing procedures

Based on the pilot data (**Supplemental Figure A**), we chose 300 µM moxifloxacin HCl, 300 µM praziquantel, 300 µM amodiaquine HCl, and 200 µM TAF. We also tested TAF mixed with 500 µM 6-MF and TAF mixed with 2.5 mM sucralose to understand their taste-modifying properties (***Table 1***).

All testing was conducted remotely using sensory software (RedJade Sensory Solutions, Martinez, CA, USA). To assess the perceived intensity of the stimuli, we asked participants to use a generalized visual analog scale (gVAS) [37, 38], due to its ease of interpretation [38] and to enable valid comparisons across groups with different sensorial experiences [37, 39].

Participants were instructed to rate the bitterness intensity of the solution (horizontal line scale ranging from “0—No Bitterness” to “100—Strongest Imaginable Bitterness”). To minimize additional variability in scale usage across individuals—since the same word might mean something different to different people—participants received a 10-min <u>training session</u>. They were first provided with verbal and written instructions on how to use the scale and practiced with graphical examples based on their food experience (drinking water, sugar, and black coffee). Next, to acclimate participants to the intensity of samples, they underwent a brief <u>warm-up session</u>, with three solutions, in a fixed order: (a) the diluent (2.5% ethanol, 0.5% Tween-80) used for 6-MF; (b) sodium chloride (256 mM), a positive control as a taste quality check; and (c) TAF (200 µM), a bitter reference for test-retest reliability with which we have previous experience [30]. Participants were asked to focus on the bitterness intensity and separate how intense a sample was from how much they liked or disliked it. All samples had an interstimulus wait time of 30 seconds, during which participants were instructed to rinse *ad libitum* with water before proceeding to the next sample, to mitigate any lingering taste effects.

Once participants were familiar with the tasting experience and how to use the gVAS, they rated the remaining nine stimuli: PROP, moxifloxacin HCl, praziquantel, amodiaquine HCl, TAF, TAF + 6-MF, TAF + sucralose, sucralose, and sodium chloride. Presentation orders of the stimuli were randomized and counterbalanced across all participants to mitigate order effects.

To characterize the diversity of the sample, after participants completed the taste test, they were asked to self-report their ancestry: city, state, and country of their mother’s, father’s, and grandparents’ birthplaces (when known). Our original intention was to create several categories within the continents (e.g., East and West Africa), but this plan was not practical, mostly because genetic analysis showed that many people fell between categories or were otherwise hard to classify.

#### Special procedures in place for COVID-19

This project took place during the COVID-19 pandemic, and all sensory testing (except for pilot testing to establish the concentration range) was conducted remotely. The taste and saliva collection kits were mailed to participants, and guided testing sessions were conducted via the video conference platform Zoom. Panelists and participants were screened for symptoms of COVID-19 on an emergency basis, which is permitted both by the Common Rule (Federal Policy for the Protection of Human Subjects, 38 CFR §16.108(a)(3)(iii)) and by FDA regulations (21 CFR §56.108(a)(4)).

### 2.5. Saliva collection, DNA purification, and genotyping

After sensory testing was completed, participants expectorated into an Oragene DISCOVER OGR-600 saliva collection tube (DNA Genotek, Ontario, Canada) with chemicals to preserve genomic DNA, for later extraction and purification, and mailed the tube to our laboratory in a prepaid stamped envelope. The genomic DNA was extracted, purified, and quantitated using the manufacturer’s protocol (DNA Genotek, Ontario, Canada) and diluted to 10 ng/ml for genotyping of *TAS2R38* bitter taste receptor SNPs (rs10246939, rs1726866, and rs713598). Additionally, the genomic DNA samples were shipped in four batches to the Center for Inherited Disease Research (CIDR), where they were genotyped using the Global Diversity Array (Illumina, San Diego, CA, USA) per the manufacturer’s procedures. The genetic data were cleaned using the CIDR analysis quality control pipeline.

### 2.6. Data analysis

We evaluated the results from pilot testing, collected the sensory data from participants, cleaned these data, computed a bitter suppression score to evaluate the effectiveness of the blockers, and determined whether bitterness ratings differed by age and sex. Participants were classified by ancestry using self-reported and genetic data, and we then evaluated the bitterness of medicines and suppression effects of blockers by ancestry. Correlations among the bitter ratings of medicines were examined for pairs that were similar or different in their degree of bitterness. We conducted a genome-wide association analysis to find genetic associations with person-to-person differences in bitterness perception. All statistical analyses were performed using R 3.6.0 (37) and RStudio 1.2.1564 [40]. The R scripts and compiled data used for this analysis are available on GitHub (https://github.com/MonellReedLab/Gates).

#### Calculation of bitter suppression scores

We tracked participants from recruitment and consent through sensory testing and saliva collection steps and removed participants who were ill (e.g., cough via self-report) on the day of sensory testing (***Figure 1***). It is not possible to evaluate the potency of a bitter blocker for participants who are insensitive to the bitter taste of a particular stimulus. Therefore, to evaluate the effectiveness of bitter blockers on TAF, we removed participants who experienced no bitter taste from TAF (with ratings ≤25 on the 100-point gVAS) and created a bitter suppression score by subtracting the bitterness intensity rating of the mixture of TAF plus 6-MF or sucralose from that of TAF alone, presented as percentage bitterness vs. TAF alone. Larger suppression scores mean more bitter suppression (and better bitter blocking). <u>All statistical analyses were performed on all bitter ratings</u> except for the bitter suppression scores.

**Figure 1.**
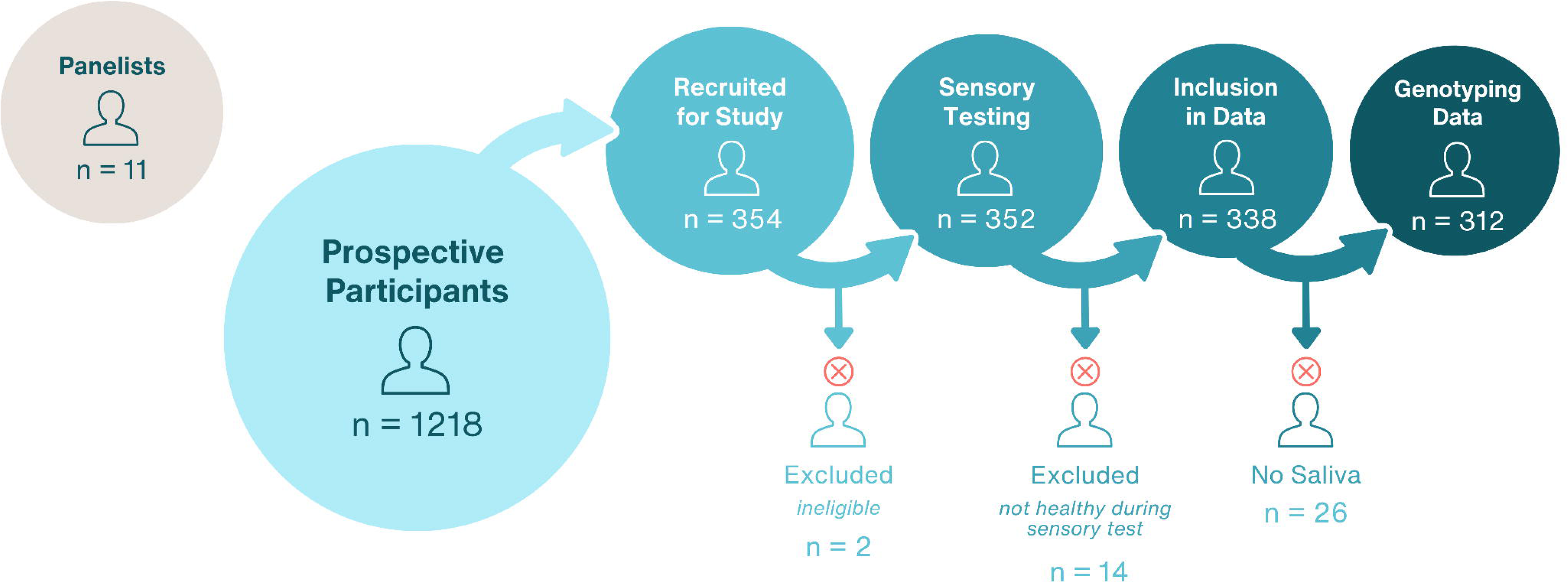
Numbers of panelists and participants in the pilot and large-scale studies.

#### Bitter taste intensity and effects of age and sex

In general, sensory ratings for some taste stimuli are more affected by age than others [41]. Therefore, we first examined the effects of age and sex on the sensory ratings to decide whether to include these variables in subsequent analyses. We conducted a Pearson’s correlation efficient for age and the bitterness rating for each of the nine sensory stimuli (PROP, moxifloxacin HCl, praziquantel, amodiaquine HCl, TAF, TAF + 6-MF, TAF + sucralose, sucralose, and sodium chloride) and tested whether correlation coefficients were statistically different than zero. We assigned sex at birth by evaluating the genetic markers on the X chromosome and tested whether differences in bitterness ratings depended on chromosomal sex using a Wilcoxon rank sum test (26 subjects with no available genotype data were excluded from this analysis).

#### Bitter taste intensity and ancestry

We grouped people by ancestry using parental and grandparental birthplace and genotype to generate a harmonized, best-available classification (see “*Harmonizing ancestry*,” below) and compared groups by ancestry using a general linear model that included age and sex as covariates. Specific group differences were evaluated using Tukey post-hoc testing.

#### Efficacy of bitter blockers and ancestry

We evaluated the bitter suppression scores for 6-MF and sucralose by ancestry groups using general linear models similar to the taste intensity analysis.

#### Similarity and differences between pairs of taste stimuli

We used a Pearson’s correlation test statistic and displayed the results in a heat map with Pearson correlation coefficients. We also compared pairs of correlations to determine if some medicines were more similar than others.

#### Genotype processing, sex, imputation, and mapping

CIDR provided genotypes of ∼1.8M variants from 312 participants after typing with the Global Diversity Array (8V1-0); 96% of the variants were autosomal, 3.9% were from the sex chromosomes, and 0.1% were mitochondrial. The missing rate for genotyping ranged from 0.27% to 0.43% for the four genotyping batches. Using the Global Diversity Array assay pipeline, CIDR assigned the biological sex of each participant. Ten participants were not classified using this method; for these we used the sex-assignment function implemented in *plink2* [42]. To ensure accurate imputation of additional variants, we dropped variants from sex chromosomes, those with a minor allele frequency <5%, and a genotype call rate <90%, as well as all insertion and deletion variants and variants with strand ambiguous (A/T and C/G) calls. Thus, we used 782,080 autosomal variants as input into the Michigan Imputation Server [43] using the reference genome Haplotype Reference Consortium release 1.1. After imputation, we filtered out variants that (a) had a low minor allele frequency (<5%), (b) were not in Hardy-Weinberg disequilibrium (using a threshold p-value of <1e-6), (c) had a genotype call rate <0.9, or (d) had an imputation score <0.3. The remaining 6,340,313 variants were used in the downstream genetic analyses. The variant map location reported is relative to GRCh37/hg19.

#### Harmonizing ancestry using four methods

Participants self-reported the birthplace of their parents and grandparents (if known), and we compared this information with three genotype classifications of ancestry. We used multiple methods, each with different strengths and weaknesses, to make the best available classification. (1) We used all available genotype data and computed the genetic distances to the genomes of 1,092 participants in the Phase 1 analysis of the 1000 Genomes Project [44] using functions implemented in the *GRAF-pop* program ([45]; version 2.0). (2) Using a model-based method with 2,538 Ancestry Informative Markers, we took a classic Bayesian approach to assign individuals to populations (*STRUCTURE*; version 2.3.4) [46]. (3) A principal components classification method was implemented in *plink 2.0* [42]. (4) For 26 participants with no genotype data, we used their self-reported ancestry.

#### Genome-wide association

We conducted a mixed linear model (MLM)-based genome-wide association analysis implemented in *GCTA* (version 1.94.1) [47] using 6,340,313 variants on the 22 autosomes with genetic relationship matrix (GRM) estimated individual pairs from genome-wide SNPs. Ten principal components (eigenvec) were used to correct the ancestry structure, and age and sex were included as covariates. We assumed a genome-wide significance threshold of p<5e-8 and a suggestive threshold of p<1e-5. For significant associations, we plotted results using the web tool LocusZoom [48]. We merged the GCTA output, which included test statistics for each variant and its genomic location, with variants annotated using GRCh37.p13 [49].

## 3. Results

### 3.1. Participant screening and demographics

From >1,000 potential participants screened, we studied fewer than one-third (***Figure 1***). This study took place at the height of the SARS-CoV-2 pandemic and required a change in protocol from a standard sensory testing because of the quarantine procedures in effect: all participants were shipped the chemosensory stimuli and were tested in their homes (or a similar location) via Zoom. Taste loss is a symptom of COVID [50], so all participants were screened for COVID by questionnaire; although none reported having it on the day of sensory testing, out of an abundance of caution some participants were excluded for any signs of even the most minor illness (N=14). We used an unusually large number of recruitment methods, with some more effective than others in identifying eligible participants (e.g., ***Supplemental Figure B***). In total, we used the data from 338 participants. This includes sensory data collected from 26 people who did not provide a saliva sample; their data were not included in statistical analyses relying on genotype, such as sex effects, genome-wide association, and PROP bitterness by *TAS2R38* genotypes. Participants were adults ranging from 18 (the minimum eligible age) to 73 years old (average = 32 years; ***Table 2***) and were 62.7% (n = 212) female as measured by sex chromosome genotype, 29.6% (n=100) male by genotype, and 7.7% (n=26) missing genotype data. No attempt was made to classify participants by gender with a self-reported question.

**Table 2.** Participant characteristics.

|  |  |  |
| --- | --- | --- |
| **Total participants (N)** |  | **338** |
| **Age, mean ± SD, years old** |  | **32.5 ± 11.5** |
| **Sex, N (%)** |  |  |
| Female |  | 212 (62.7) |
| Male |  | 100 (29.6) |
| Not predicted<sup>a</sup> |  | 26 (7.7) |
| **Ancestry harmonized for data analyses, N (%)** |  |  |
| European |  | 57 (16.9) |
| African |  | 127 (37.6) |
| Asian |  | 81 (24.0) |
| South Asian |  | 73 (21.6) |

<sup>a</sup>No DNA for sex chromosome determination.

### 3.2. Age and sex analysis

For all the medicines tested, younger and older adults were similar in their ratings of bitterness intensity. Older people rated sucralose and sodium chloride as slightly more bitter than did younger participants, but these effects were small and influenced by only a few people, especially for sucralose (and only a few people reported sucralose as bitter; ***Figure 2***). Genetic females (participants with female sex chromosomes) rated three of the medicines as more bitter than did genetic males: moxifloxacin, amodiaquine and TAF (***Supplemental Table D***). This sex effect was strongest for moxifloxacin (rated 57% more bitter by females) and weaker for amodiaquine (22%) and TAF (15%). To account for these effects, age and sex were included in subsequent analysis as covariates.

**Figure 2.**
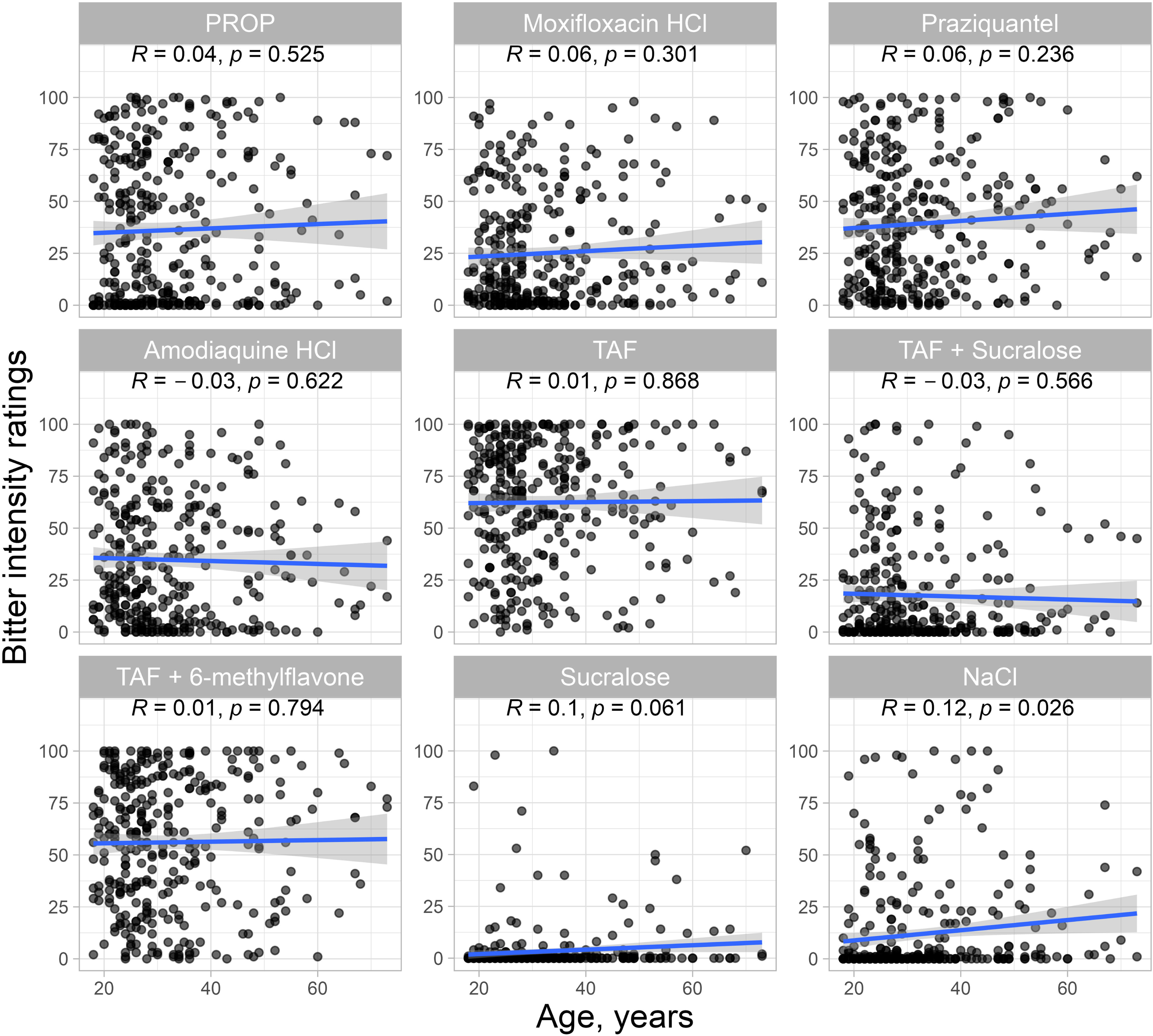
Correlation between age and bitter intensity ratings for nine sensory stimuli (N=338), with Pearson’s correlation coefficients (R).

### 3.3. Harmonizing self-report and genotype to assign ancestry

The ancestry groups are European, African, Asian, or South Asian. We classified people into these ancestry groups based on their self-reported birthplace of their parents and grandparents, their self-reported ancestry, and the ancestry groups suggested from the three types of genetic analysis mentioned above. **Supplemental Table E** shows the classification of each participant, and the results of the principal components classification method are presented in **Supplemental Figure C.**

### 3.4. Ancestry effects on bitterness ratings and bitter suppression

We tested for differences in the bitterness intensity ratings by ancestry (***Figure 3***, ***Table 3***). Large person-to-person differences in the ratings of bitterness were apparent for all medicines and in all ancestry groups. Within each group, some people rated the medicine at the top of the scale, indicating intense bitterness, whereas others rated it as weak as water. Despite this wide range in responses, group averages of bitterness ratings of PROP and amodiaquine did statistically differ by ancestry. On average, PROP was more bitter for people of Asian ancestry compared with other ancestries, and amodiaquine was more bitter for people of European than for people of African ancestry.

**Table 3.** Bitter intensity ratings of five drugs (PROP, moxifloxacin, praziquantel, amodiaquine, and TAF), TAF plus a bitter blocker (sucralose or 6-MF), and two control solutions (sucralose, sodium chloride) for all participants and for the four ancestry groups.

| Tastant | N | Bitter Rating [Mean (SD)] |  |  |  |  |
| --- | --- | --- | --- | --- | --- | --- |
|  |  | Overall (N = 338) | European (N = 57) | African (N = 127) | Asian (N = 81) | South Asian (N = 73) |
| **TAF** | 338 | 62.4 (29.2) | 64.0 (26.2) | 63.1 (30.2) | 60.0 (30.0) | 62.5 (29.5) |
| **Moxifloxacin HCl** | 338 | 25.1 (26.8) | 23.4 (25.5) | 27.3 (28.6) | 23.3 (26.1) | 24.4 (25.3) |
| **Praziquantel** | 338 | 39.3 (30.3) | 37.9 (26.5) | 39.1 (31.9) | 41.6 (30.1) | 38.2 (30.9) |
| **Amodiaquine HCl** | 338 | 34.7 (30.0) | 42.9<sup>a</sup> (29.1) | 31.3<sup>b</sup> (29.2) | 34.2<sup>a,b</sup> (30.0) | 34.8<sup>a,b</sup> (31.5) |
| **PROP** | 338 | 36.2 (34.4) | 32.5<sup>a</sup> (36.9) | 27.8<sup>a</sup> (32.1) | 55.2<sup>b</sup> (31.0) | 32.7<sup>a</sup> (32.4) |
| **TAF + sucralose** | 338 | 17.6 (25.4) | 21.6<sup>a,b</sup> (24.2) | 12.4<sup>a</sup> (22.8) | 23.7<sup>b</sup> (28.7) | 16.7<sup>a,b</sup> (25.4) |
| **TAF + 6-MF** | 338 | 56.1 (31.1) | 58.8 (28.0) | 59.2 (31.6) | 53.9 (31.9) | 50.9 (31.5) |
| **Sucralose** | 338 | 3.3 (12.1) | 3.7 (9.8) | 3.1 (12.6) | 1.9 (7.2) | 5.0 (16.3) |
| **Sodium chloride** | 338 | 11.9 (23.3) | 9.3 (18.2) | 12.7 (23.4) | 7.2 (19.8) | 17.8 (28.5) |

<sup>a,b</sup> Significant difference ( $P \leq 0.05$ ) from linear models with age and sex as covariates, by Tukey post hoc test.

**Figure 3.**
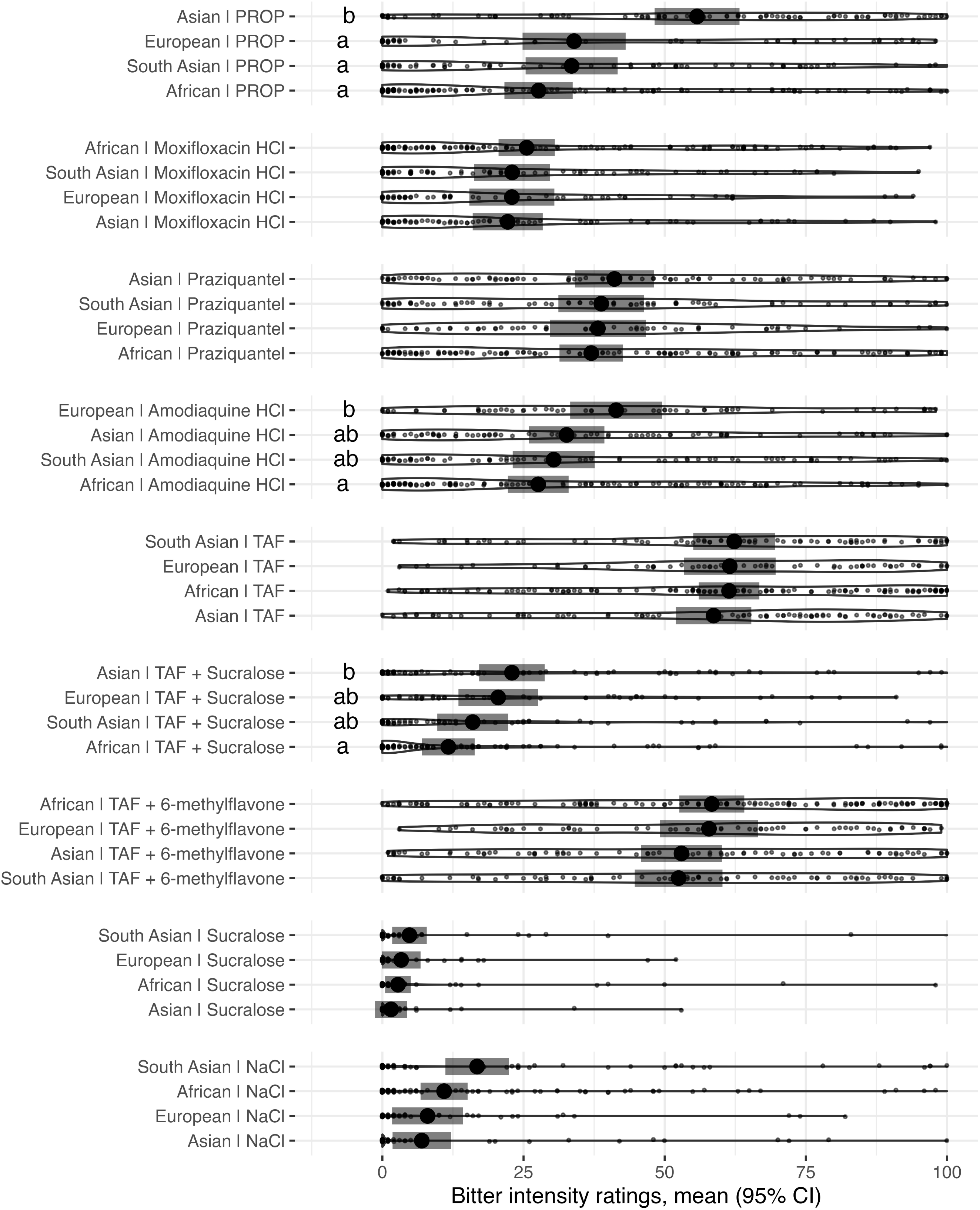
Bitter intensity ratings of five drugs, two mixtures of TAF plus a bitter blocker, and two control solutions, by people of different ancestries (N=338). Points represent individual data; violin plots overlay with the widest outline indicating the greatest number of responses; black dots and gray bars depict means and 95% confidence intervals based on linear models with age and sex as covariates. ^a, b^ Significant differences (P ≤ 0.05) between ancestry groups with Tukey post hoc tests. For details, see ***Table 3***.

Across all participants, the median bitterness suppression was best for sucralose (91%) and worst for 6-MF (8%). There were ancestry differences in how bitter TAF was when sucralose was added (***Figure 3***; an analysis including all participants) and in the suppression score for sucralose or 6-MF (***Figure 4***; computed only participants who reported TAF was bitter).

**Figure 4.**
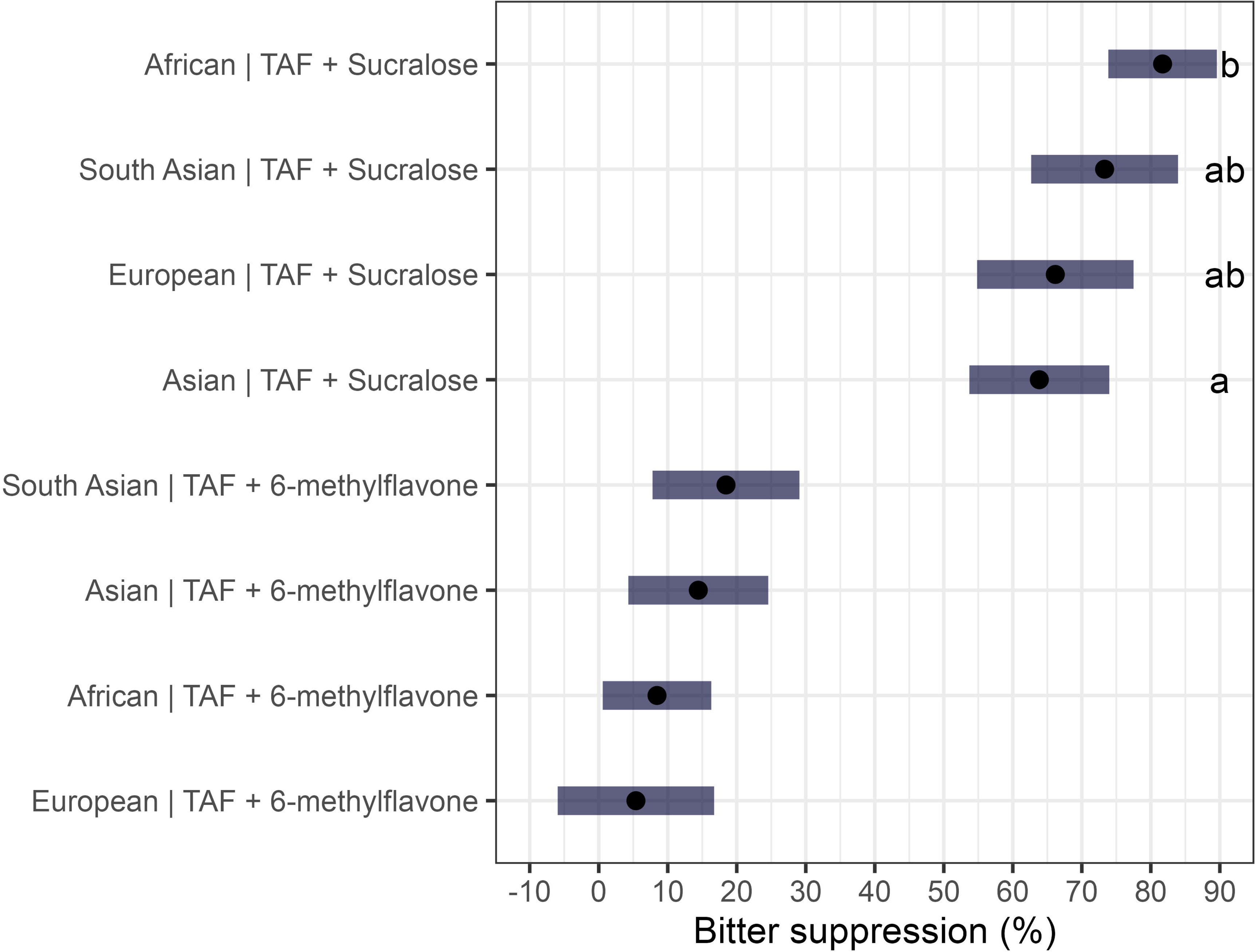
Bitter suppression (%) of TAF by bitter blockers among participants who perceived TAF as bitter (score > 25 on the 100-point gVAS) (N=284). Black dots and gray bars depict the means and 95% confidence intervals computed based on linear models with age and sex as covariates. ^a, b^ Significant differences (P ≤ 0.05) between ancestry groups with Tukey post hoc tests.

Sucralose worked best for those of African ancestry, with some individual suppression scores approached a perfect 100%, and most poorly for those of Asian ancestry. 6-MF was less effective than sucralose but suppressed bitterness (at least somewhat) for those of South Asian, Asian, and African but not European ancestry.

### 3.5. Correlational analysis

We compared the bitterness ratings of all taste solutions and mixtures. Most correlations were significant (coefficient values 0.10-0.58) (***Figure 5***), but the most informative comparisons come from the pattern, the differences among the positive correlations among all pairs of correlations, to determine if some medicines were significantly more similar than others (***Supplemental Table F***). This comparison revealed no evidence for the concept of general “supertasters,” people who experience <u>all</u> taste stimuli as more intense than others. For example, there was no relationship between ratings of bitterness for sodium chloride and for any of the other solutions tested. The bitter medicine PROP was least like the other medicines, with a coefficient value range of 0.15-0.29 when paired with any other solution tested, whereas the pair moxifloxacin-- amodiaquine had a high correlation (r=0.54). These comparisons suggest that there may be more shared underlying mechanisms for the bitter perception among moxifloxacin, amodiaquine, TAF, and praziquantel, but less common mechanisms between these medicines and PROP.

**Figure 5.**
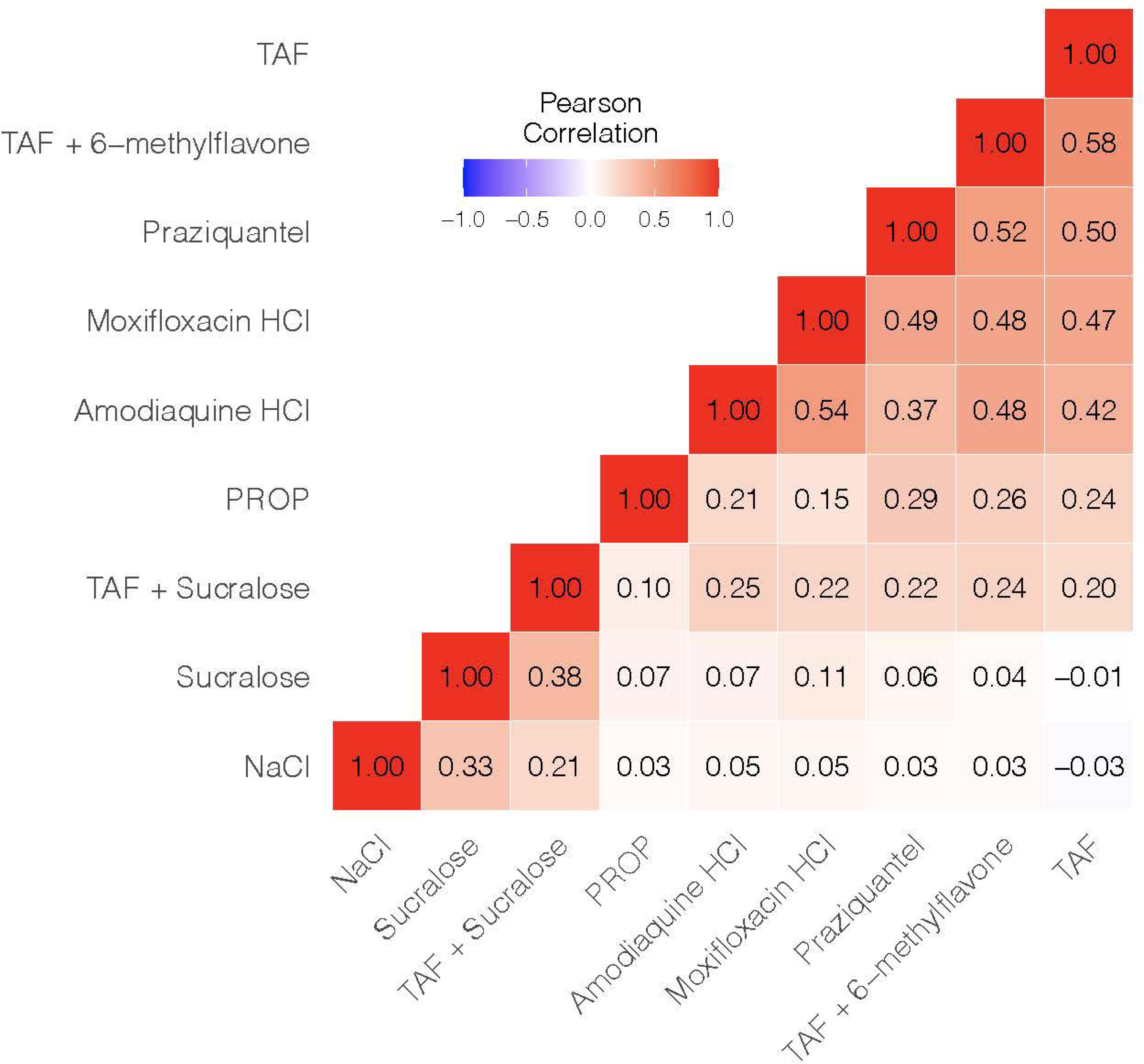
Pearson correlation in bitterness ratings among five drugs (PROP, moxifloxacin, praziquantel, amodiaquine, and TAF), two mixtures of TAF plus a bitter blocker (sucralose or 6- MF), and two control solutions (sucralose, sodium chloride). Color intensity, from blue to red, indicates the increasing correlation coefficient of the bitterness intensity ratings reported in each box.

### 3.6. Genome-wide association and all SNP-based heritability

We found significant genotype-associations with bitterness ratings for PROP for three variants in the bitter receptor *TAS2R38* (**Figure 6**), and for sucralose in the genes *RIMS2* and *THSD4* (**Figure 7**). No other taste traits had a genome-wide association that met the significance threshold. (For all association results, see **Supplemental Tables G1-8** via the link for external data.) Merging variants with gene locations is imperfect because some variant locations are within two genes and therefore results in the supplemental tables should be interpreted with this point in mind.

**Figure 6.**
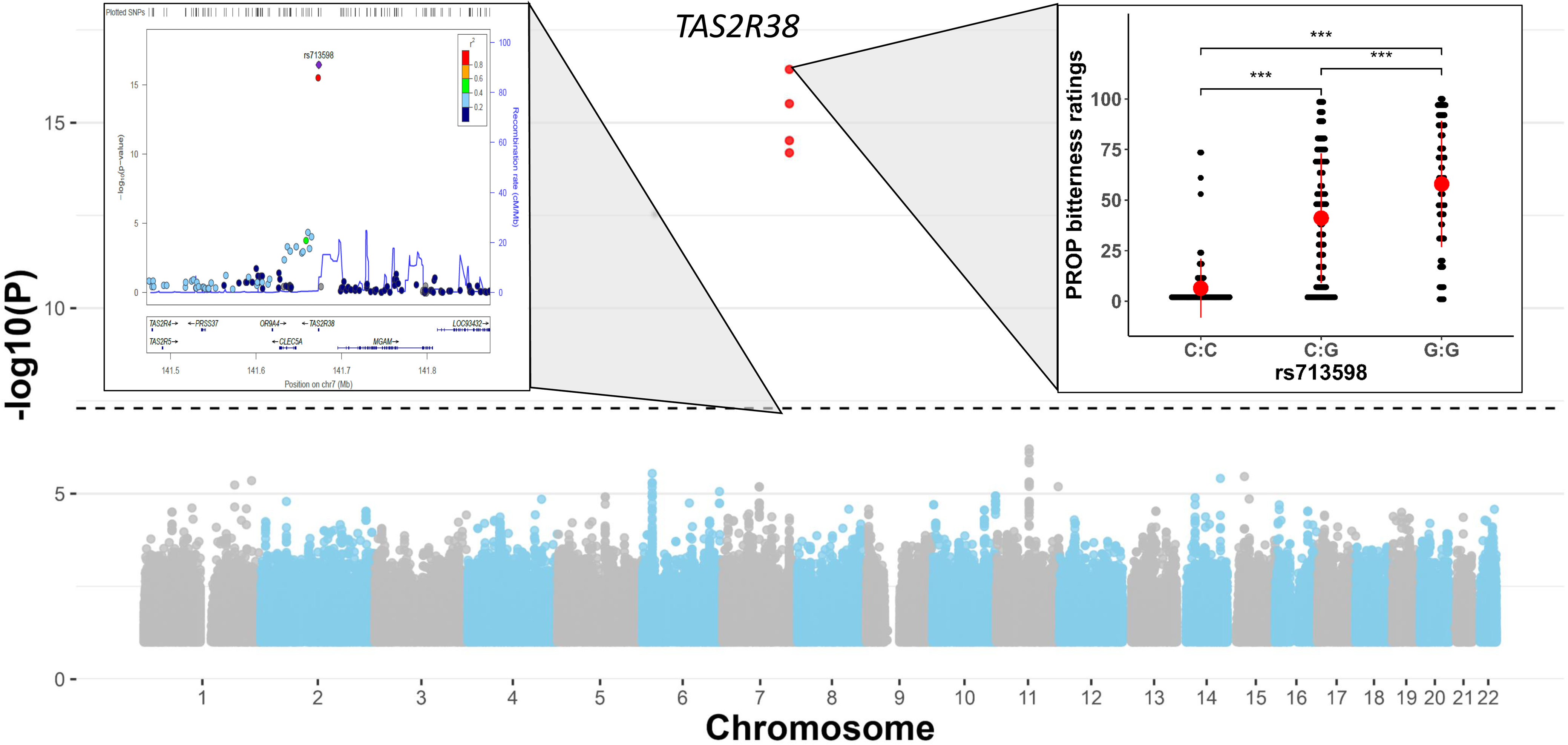
<u>Top,</u> Common variants within the bitter taste receptor gene *TAS2R38* associated with PROP bitterness ratings. <u>Top left,</u> Regional association around *TAS2R38* gene (+/- 200 Kb flanking region); color scale for the linkage disequilibrium with the top SNPs and their correlations is shown. <u>Top right,</u> PROP bitterness ratings grouped by the three rs713598 genotypes of *TAS2R38*. ***Tukey post hoc test significance with P-value ≤ 0.001. <u>Bottom</u>, The y-axis of this Manhattan plot shows the association P-value for each SNP (displayed as -log_10_ of the P-value). The black dashed line indicates the genome-wide significant threshold of P = 5.0e^-8^. Missense variant rs713598 of *TAS2R38* is the most significant SNP.

**Figure 7.**
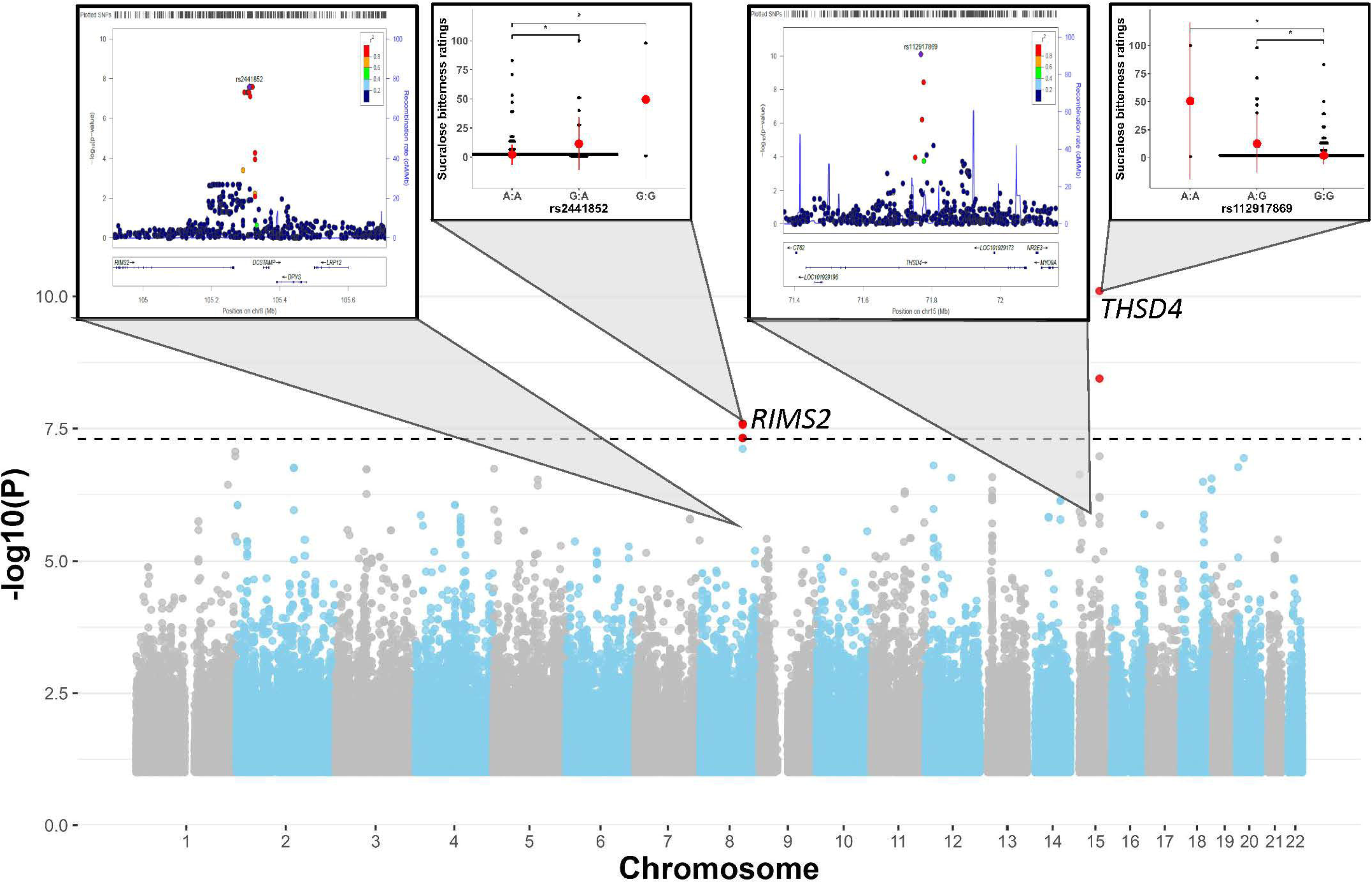
Noncoding common variants associated with sucralose bitterness ratings. rs2441850 near *RIMS2* and rs112917869 within the intron region of *THSD4* reach genome-wide significance. For other details, see Figure 6. *P-value ≤ 0.05.

### 3.7. Data reliability

Data reliability was confirmed by the ability to distinguish the bitterness of slightly bitter (e.g., diluent - data not shown, sodium chloride, and sucralose) from moderately to highly bitter solutions (TAF and other drug solutions) (***Figure 3*** and ***Table 3***) and the significant correlations between the warm-up and test samples (e.g., R=0.69 for TAF and R=0.75 for sodium chloride). The expected relationship between genetic variation in the *TAS2R38* bitter receptor and perceived PROP bitterness also confirmed both the reliability and validity of the data. For two haplotypes constructed with three common variants of the *TAS2R38* bitter receptor, AVI is the bitter-insensitive haplotype, and PAV is the bitter-sensitive haplotype. Average PROP bitterness was significantly lower for participants with the AVI/AVI diplotype (N=64) than for those with the AVI/PAV diplotype (N=119) and highest in those with the PAV/PAV diplotype (N=88).

Consistently, genome-wide association was able to identify *TAS2R38* polymorphisms (rs713598), with a strong association between the rs713598 (C:C, C:G, and G:G genotypes) and PROP bitterness ratings (for differences in PROP bitterness ratings between these three groups see ***Figure 6***).

## 4. Discussion

### 4.1. Individual differences

Our study shows that the bitterness of medicines, even those infamous for their bad taste, like moxifloxacin [51], differs strikingly among people, which may explain in part why some patients do not take the medicine as prescribed. For some, the medicines may be so bitter, and they cause nausea, choking, or vomiting [52]. One implication of these results is that, while medicine formulations may be developed to please an average person, they may not reduce the noxious taste for all (or, in some cases, can make it worse, depending on what is added). One simple step would be to pre-qualify panelists during formulation development, ensuring they are bitter sensitive to both a medicine’s active ingredients and its excipients. (The same precaution could be taken for other unpleasant flavors, like sourness and astringency, not studied here). These individual differences and their impact on the design of foods, drinks, and medicines is not new, an observation sometimes captured in the Latin phrase *de gustibus non est disputandum*: “in matters of taste, there can be no disputes,” referring to the incomparability of any two people’s tastes. These differences are part of the range of human sensory experience but present a challenge for formulating medicines that are acceptable to all.

We found large differences in the efficacy of the two taste modifiers we tested here to block the bitterness of the antiviral drug TAF: sucralose was very effective, with some participants reporting that TAF mixed with a high concentration of sucralose was not bitter at all, yet others reported that sucralose *added* to the bad taste, rate it as worse than TAF alone. Results also differed across participants for 6-MF, which was less able to reduce bitterness overall compared with sucralose. Sweeteners at high concentrations have been widely used in medicine formulations as partly effective bitter-reducing excipients [22, 53], due to their bitter-masking and potential blocking effects [54]; sucralose also had a greater bitter suppression than other flavoring agents, taste modifiers, and bitter blockers in formulations of primaquine, an important antimalarial drug [16].

### 4.2 Sources of the individual differences

To examine the origin of these individual differences in bitter taste perception, we compared participants of different age, sex, ancestry, and genotype. Genotype was an especially important factor: for some medicines, variation in a single bitter receptor can account for nearly the full range of taste responses (as with the classic example of *TAS2R38* and PROP [55]). We used this drug-receptor pair as a control in this experiment to ensure we could detect this type of genetic effect. Although we saw the expected genetic relationship when participants tasted PROP, we did not observe any other single-genotype relationships between a medication’s bitterness and any variant in a bitter receptor or any other genes. Thus, the PROP genotype-phenotype relationship – between one genotype and the bitterness of a drug — is more the exception than the rule, at least for the medications tested here.

In cases where a drug is a ligand for several bitter receptors [56], variants within each may make small addition toward bitterness perception that individually are hard to detect. An example of this polygenic effect is TAF, known from previous studies to stimulate at least four receptors [29]. Genotypes can also interact, with little observable phenotypic effects unless all the influential alleles are non-functional in a particular person – a ‘perfect storm’ that makes the person very sensitive or insensitive. For none of the other medicines we tested do we have information on which or how many bitter receptors they activate – but the correlations we observed among them suggest they may share several bitter receptors; testing these medicines with receptor-based heterologous expression assays [29] will help identify the associated receptors. Furthermore, like many studies using genetic association, larger numbers of participants will increase our ability to detect small additive or interacting effects.

These same points apply to excipients: the person-to-person variation in at least one excipient, sucralose, may be explained by genetic variation, although not within taste receptors. Two genomic regions met the conventional threshold for statistical difference in the genome-wide association analysis within *RIMS2* and *THSD4*, two genes not known to be part of the human taste signaling system. In a twin study *RIMS2* variants explained, in part, differences in liking for high-fat foods [57]; its protein product regulates synaptic membrane exocytosis. People with *RIMS2* loss-of-function variants have visual and pancreatic defects [58], and this information, coupled with our knowledge that sweet taste cells lack conventional exocytotic synapses, suggest that *RIMS2* may act to modify sucralose perception in cells other than taste receptors (e.g., in the brain or pancreas). *THSD4* variants are associated with an array of human diseases and traits that have little apparent direct effect on the sense of taste (e.g., [59]), and the mechanistic path from the function of this protein to the perception of sucralose is unclear.

Whether these genetic associations can be replicated in other populations will tell us whether mechanistic studies on these genes and their protein products are warranted.

### 4.3. Ancestry

This study was conducted in the United States and Canada, which have many recent immigrants from all over the world. We chose to study these immigrants as a proxy for geographic diversity. We studied first-, second-, or third-generation immigrants, so there was little generational time for significant admixture; and our expectation is that our sample is very similar genetically to those still living in Africa, Asia, and South Asia. Two out of the five medicines tested were rated as more bitter (on average) by people of a particular ancestry.

Moreover, we saw ancestry differences in the efficacy of the two taste modifiers in suppressing the bitterness of TAF. In medicine formulations, excipients are often used not only to affect taste but also to facilitate drug delivery, enhance stability, or improve other qualities. Ancestry differences may also contribute to individual differences in the perception of these additional excipients, hence contributing to the flavor qualities of formulations. For example, in a recent study, an over-the-counter sweetened suspension of the anti-inflammatory drug ibuprofen, commonly used for children, was perceived sweeter and less irritating (but not differing in bitterness) by people of African than of European ancestry [60]. Thus, to achieve broadly palatable medicines, formulations must be tested with people with diverse ancestry.

We used several sources of information to classify people into ancestry groups, comparing self-reported ancestry based on family history (geographic birthplace of grandparents) and genotype. However, most of our participants were cosmopolitan, with grandparents from different countries, sometimes different continents. This may have arisen in part because in recruitment we tapped professional networks with a focus on academic environments. We supplemented this family history with ancestry classification based on genotype, using several statistical methods that converged to similar categories. While we are confident of the classification to the broad regions used in this study (e.g., African, Asian), finer-grained studies of geographic populations or of people with a particular ancestry are best conducted in the home country among groups with low rates of migration and immigration.

### 4.4. Value of human diversity in taste studies

Studies that include people primarily of one ancestry, often European, can have serious limitations [61], captured by the phrase “missing diversity”. Including participants from Africa, Asia, and other regions, as well as Europe, can increase the generalizability of results. It also has the potential to uncover new information. For example, by including people of African ancestry, we established that sucralose can be a powerful reducer of bitterness, an effect less apparent in people of other ancestries. For taste studies, especially those that depend on bitter taste receptor genes, it is especially important to include people of diverse ancestry because taste receptors, which change over evolutionary time [62, 63], show large differences in allele frequencies across human populations. For example, the *TAS2R38* alleles that contribute to the sensitivity to PROP bitterness are present at higher rates in South Asian populations [20], which results in differences in ratings of PROP bitterness across ancestry groups. Including people of diverse ancestry was essential to the discoveries made here. Some drugs work better in some people than others, partly because of genotype and ancestry [64]. This body of knowledge, known as pharmacogenetics, is starting to provide more effective treatments to people worldwide. Here, we expand this perspective to medicine formulation, using this knowledge to create effective medicines that people will take as prescribed.

### 4.5. Age and sex effects

We expected that ratings of bitterness for the medications might be higher and the efficacy of excipients lower among younger compared with older participants, but we found little or no effect of age on reported bitterness. Children rate medicines as more bitter compared with adults, and known bitter-reducing excipients tend to be less effective for them [23]. Although we studied only adults, the age range was large (18-73), yet we did not detect lingering taste effects of development in our younger participants. We were also surprised that the relationship between age and reported bitterness was the same for all bitter medicines, because another study showed differences among bitter taste solutions with age [65]. However, we may have tested medicines that taste bitter throughout the lifespan.

We expected women to rate bitter taste stimuli as more intense than did males, but that was true here only for some medicines. This result may be a statistical fluke, or some aspect of sex may affect the sense of taste differently for some bitter medicines than for others. Additional studies are needed to replicate these results and test for potential mechanisms for this effect (e.g., effects of sex hormones on taste receptors). We used only chromosomal sex here and did not ask about gender, which may have categorized participants differently than other studies, reducing sex differences for some medicines and exacerbating them in others (e.g., amodiaquine).

### 4.6. Sensory methods

We had participants rate the relative intensity of bitter medicines using the generalized visual analog scale (gVAS). We considered using the European Pharmacopoeia method [66], which expresses the sensitivity to a medicine relative to quinine HCl (a common bitter stimulus used in human taste studies). However, because the perception of quinine also varies by genotype [17, 35, 67], this method was not appropriate. Although the generalized labeled magnitude scale (gLMS) uses adjective labels that are better for magnitude estimation than the gVAS, thus capturing both minor and major sensory differences, gLMS data often show categorical behavior (i.e., clustering near the labels), leading to non-normal data, which can be challenging to interpret [37-39]. Moreover, the quasi-logarithmic nature of the scale confuses participants who are not familiar with it; thus, it is often used with trained or experienced participants. In this study, to test people with diverse ancestry and different sensorial experiences, we selected gVAS due to its ease of interpretation and valid comparisons across participant groups [38].

Although this scale sacrifices semantics about the magnitude of perception, it discriminates between stimuli samples as well as the gLMS does [38]. To reduce the individual variation in how people used the scale and eliminate the central tendency, we first familiarized participants with the gVAS and stimuli, by performing a training on scale use followed by a brief warm-up session with three taste samples. The strong correlation in bitter ratings between the warm-up and test ratings for TAF indicated high test-retest reliability. Moreover, we found a well-known genotype-phenotype relationship [18] between *TAS2R38* genotype and bitter ratings of PROP, validating our sensory testing method. This suggests that the testing procedure used here has potential for future research with diverse participant groups in similar experimental designs.

### 4.7. Clinical significance

How individuals perceive bitterness can significantly affect their health outcomes, especially for children and older individuals. Our findings highlight the importance of developing palatable medication that aligns with a person’s preferences. Understanding how the taste of drugs impacts medication adherence and awareness of these genetic differences allows prescribing clinicians to counsel patients about taste-related side effects, setting realistic expectations, and providing strategies to manage the symptoms.

## 5. Conclusions

The perception of the bitterness of medicines greatly differs from person to person, which can be partly explained by genetic variation. However, specific genetic contributions might be polygenic and therefore hard to detect – the strong relationship of PROP bitterness and a single genotype (i.e., *TAS2R38* variants) may be rare. The contribution of ancestry difference was common (but not universal) in our sample of recent US and Canada immigrants from Africa, Asia, and South Asia; bitterness ratings differed by ancestry for two of the five drugs. We found the same results for the perception of bitter-reducing excipients and their efficacy, which for some can contribute flavor qualities to medicine formulations that are less palatable than the drug alone. Thus, to obtain broadly palatable medicines, formulations need to be tested with people of different populations and ancestries, and with people who are known to be sensitive to the off flavors in all the medicine’s ingredients. Further study of a particular medicine with receptor-based assays can help us identify not only associated receptors but also genetics underlying its bitterness, supporting the discovery of its potential bitter blockers. Additionally, human testing with a larger number of participants can increase the ability to detect small or interacting effects of genetics and other participant characteristics, such as age and gender.

## CRediT author contributions

**Ha Nguyen:** Investigation, Data curation, Formal analysis, Visualization, Writing - original draft; **Cailu Lin:** Data curation, Formal analysis, Visualization, Writing - review & editing; **Katherine Bell & Amy Huang**: Investigation, Methodology, Resources; **Mackenzie Hannum**: Investigation, Methodology, Writing - review & editing; **Vicente Ramirez**: Investigation, Writing - review & editing; **Carol Christensen**: Conceptualization, Funding acquisition, Writing - review & editing; **Nancy E. Rawson**: Conceptualization, Funding acquisition; **Lauren Colquitt**: Investigation; **Paul Domanico**: Conceptualization, Writing - review & editing; **Ivona Sasimovich**: Investigation; Methodology**; Riley Herriman**: Investigation; Methodology, Writing - review & editing; **Paule Joseph**: Conceptualization, Methodology, Writing - review & editing; **Oghogho Braimah**: Investigation, Writing - review & editing; **Danielle R. Reed**: Conceptualization, Funding acquisition; Investigation, Methodology, Project administration, Resources, Supervision, Writing - original draft.

## Acknowledgments

We acknowledge the exceptional support from Valentina Parma in all aspects of this project, especially participant recruitment. We thank Kelly Catlin, Linda Lewis, Melynda Watkins, and Gary Moore for assistance in obtaining the drugs for sensory testing and information about their properties. We thank Natasha Rivers, Linda Flammer, and Paul A. S. Breslin for their assistance in preparing the drugs for sensory testing. Sarah Lipson helped with literature reviews and other aspects of the early phases of this research. We thank Peihua Jiang, Maureen O’Leary, and Erik Schwiebert for helpful discussions about mechanisms of reducing bitterness. Hanh Ngo, Mercy Okezue, Sarah Tishkoff, Patricia Frimpong, Patrice Hubert, Cleopatra Ogharadukun, Marius Baguma, Adeola Ayano, Dola Adeboye, Kolawole Falade, Maame Yaakwaah Blay Adjei, Ganiyat Olayinka, Jeanine Kriek (Sainsbury), Johnson Adejuyitan, Heidi Grimmer, Riette De Kok, James Makame, Shadrack Frimpong, Jane Bartonjo, Renée Hartig, Jenifer Trachtman, Bruce Hamaker, Riley Koch, Angela Nwaneri, Sabrina Wang, and Layo Jegede helped with participant recruitment. We acknowledge the Genetic Resources Core Facility (RRID:SCR_018669) and the kind assistance of Roxann Ashworth for genotyping services.

Note: PJ is supported by National Institute of Alcohol Abuse and Alcoholism under award number, Z01AA000135, the National Institute of Nursing Research and the Rockefeller University Heilbrunn Nurse Scholar Award, the Office of Workforce Diversity, and the Office of Workforce Diversity, National Institutes of Health Distinguished Scholar Program.

---

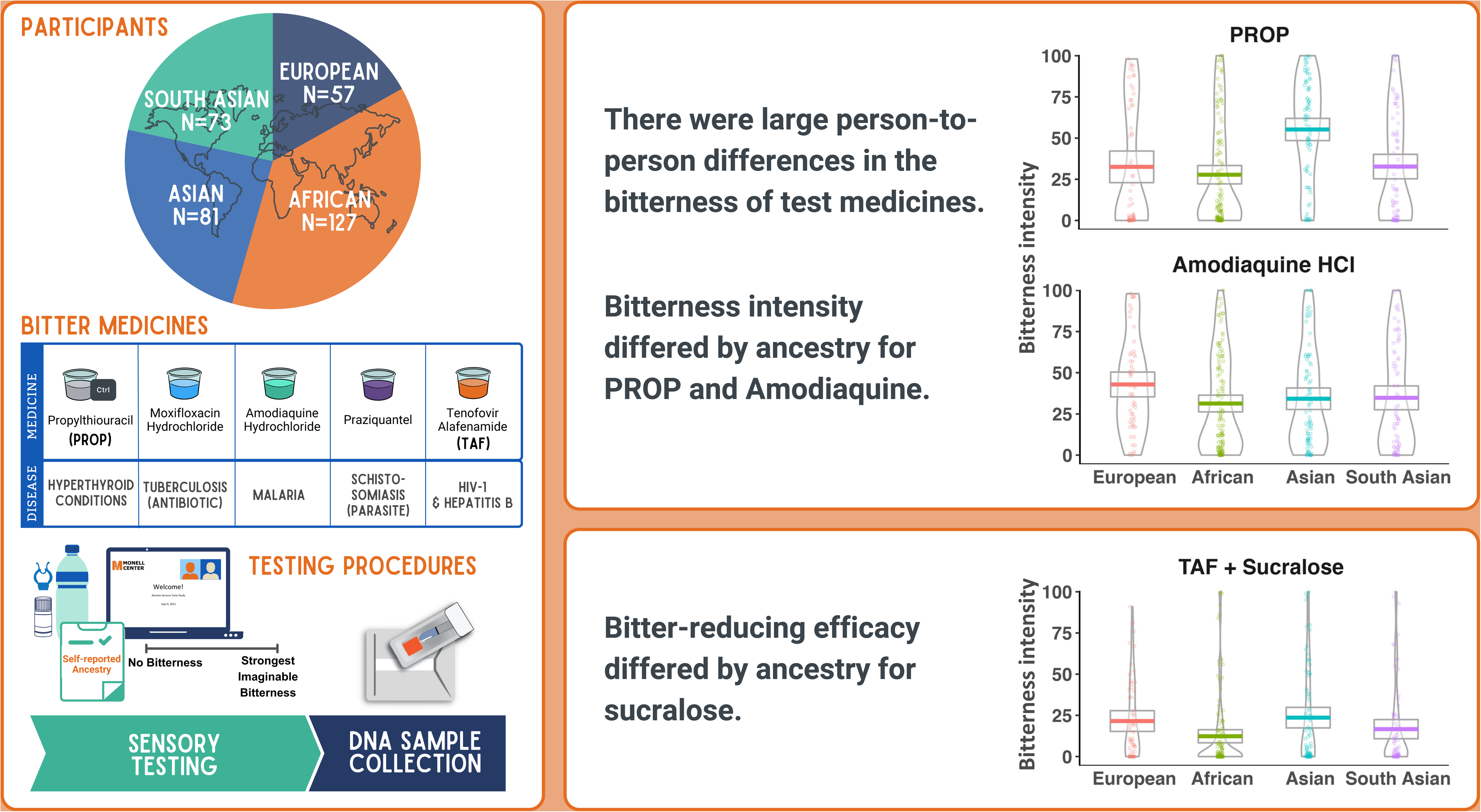

