## Supplemental Figures A-C for "Worldwide study of the taste of bitter medicines and their modifiers"

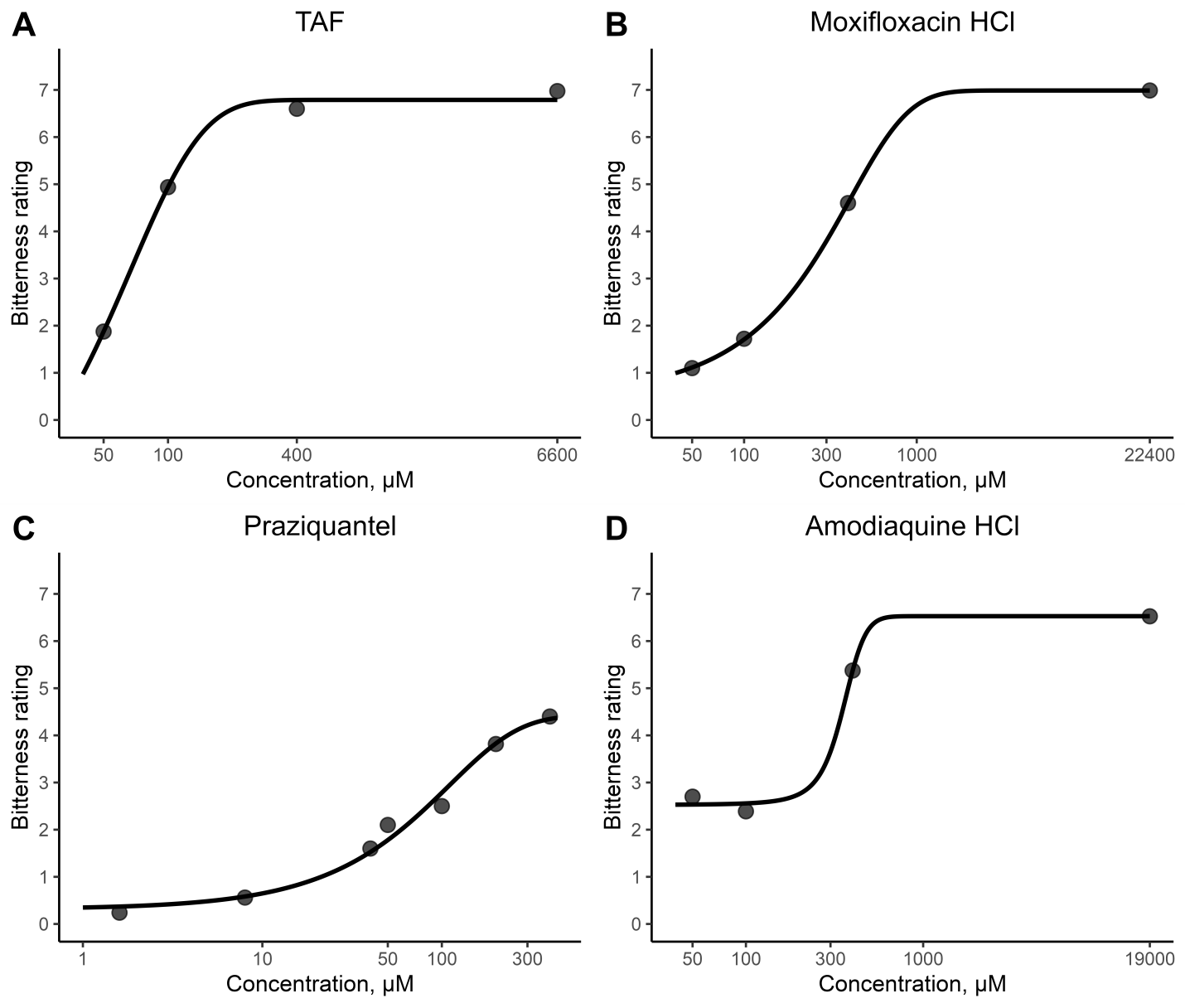
**Supplemental Figure A.** Bitterness ratings of different concentrations for (A) TAF, (B) moxiflocaxin HCl, (C) praziquantel, and (D) amodiaquine HCl

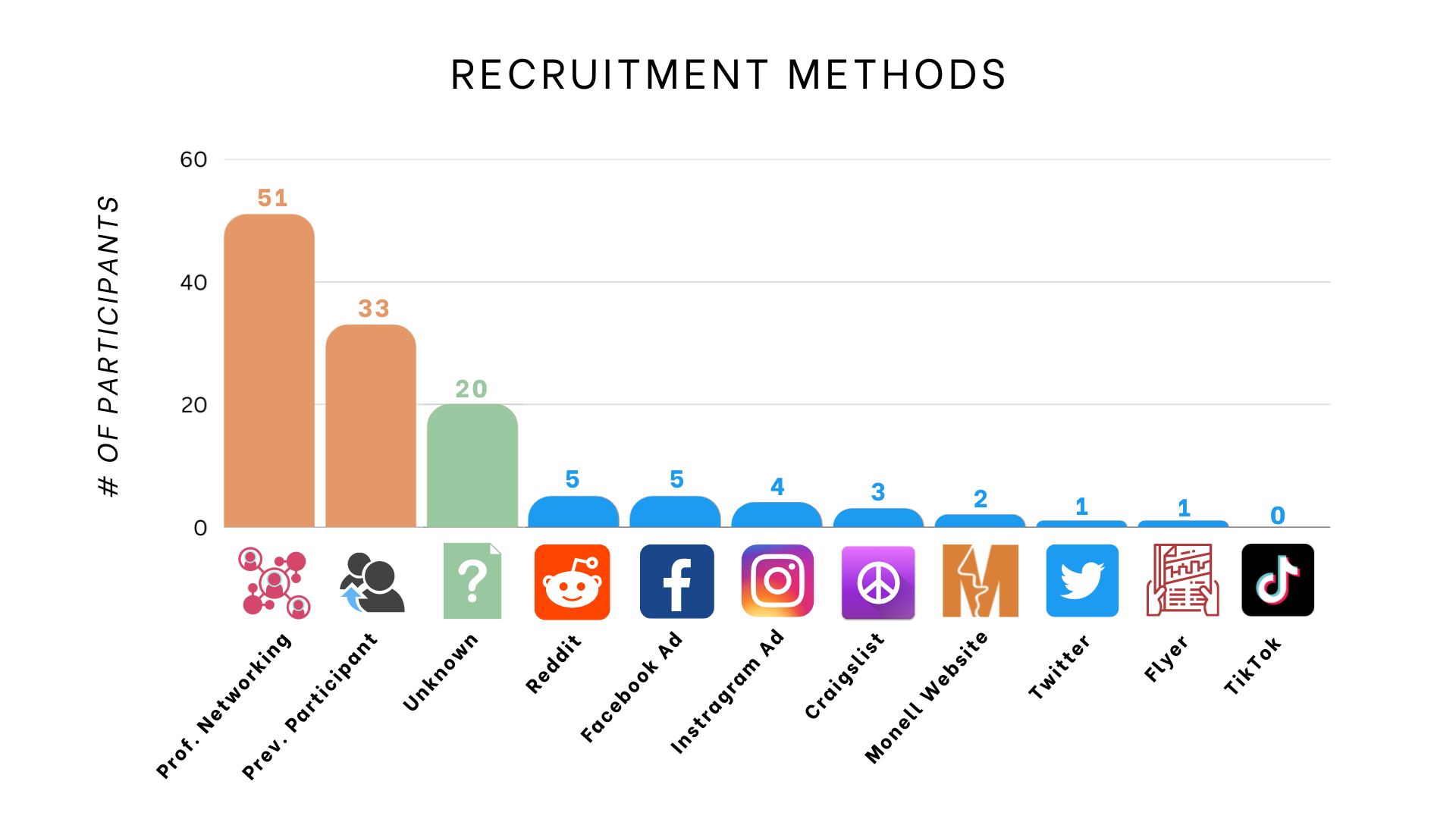

**Supplemental Figure B.** The number of African participants recruited by different methods

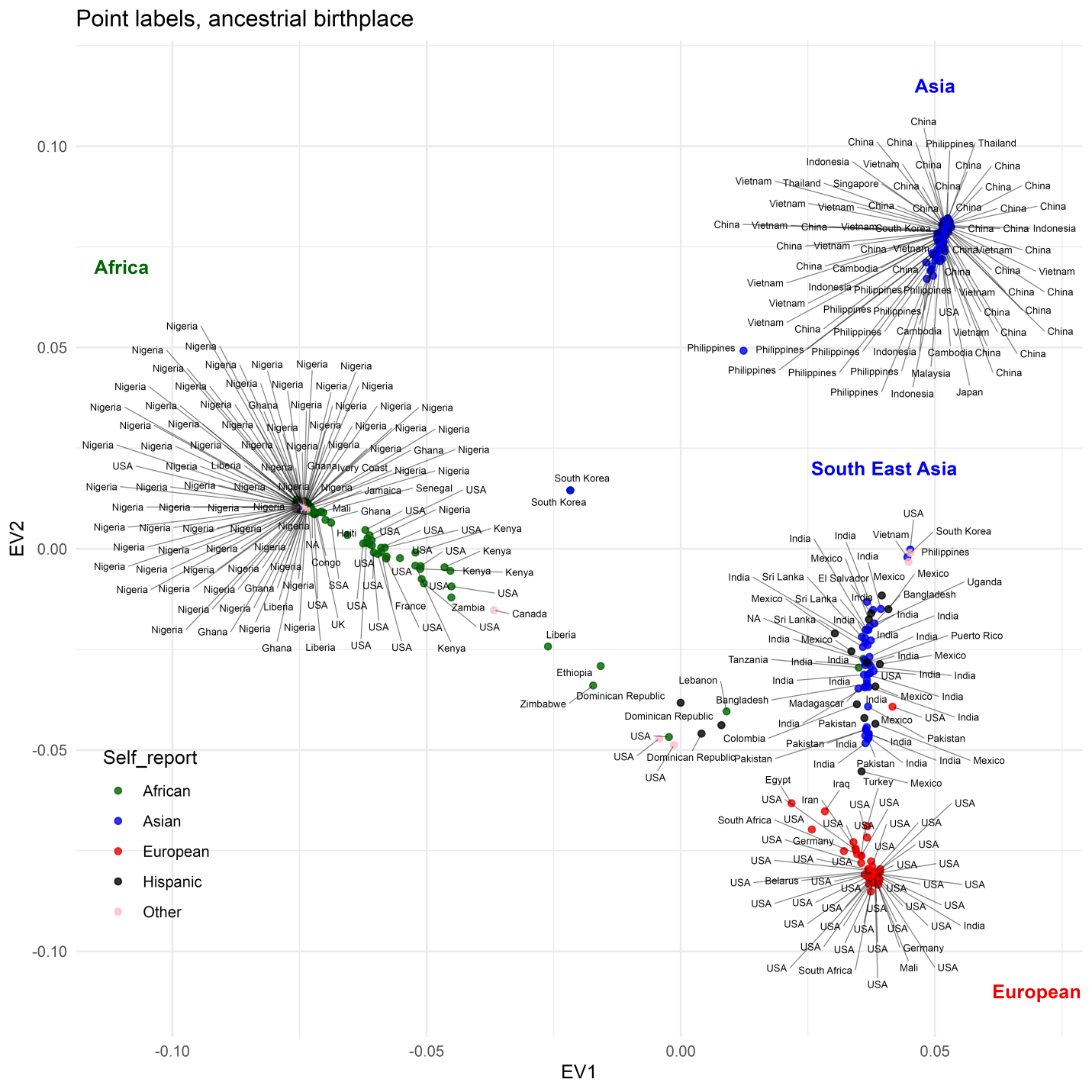

**Supplemental Figure C.** Principal components classification of participants for ancestry based on genotype and self-reported data. Point labels show ancestral birthplaces; point colors show ancestry groups according to self-reported data.
